# Predictive and Aetiological Potential of Polygenic Strata with Extreme and Moderate Disease Risks Predictions: *Case Study of Coeliac Disease GWAS*

**DOI:** 10.1101/687889

**Authors:** Adam Kowalczyk

## Abstract

We show by using example of coeliac disease (CD) that a genomic risk assessment could significantly improve efficiency of disease diagnosing. It can detect novel highly deleterious rare variants (penetrance 100%, frequency ∼1:6,700) as well as common protective variants (penetrance 0.03%, frequency ∼1:3). However, the major translational gains with potential for multi-billion-dollar cost savings in Australia or USA alone, could be in assessing patients in cohorts with moderately elevated CD risk (3% −10%) exhibiting clinical symptoms or with family history of CD. The gains result from judicious re-direction of expensive confirmatory testing towards ∼30% of the cohort with the highest likelihood of the condition (∼90% of cohort CD cases), while avoiding costs, inconvenience and risk of side-complications for the remaining majority of ∼70%.

We build our estimates using concrete results of CD Genome Wide Association Studies (GWAS) already in the public domain^1–4^. The largest of five Genomic Risk Score (GRS) models^1^ considered here deploys 228 directly genotyped Simple Nucleotide Polymorphisms (SNPs), while the simplest^2^ uses only 6 SNPs. Thus, a DNA profile supporting all these models can be easily accommodated on any commodity, Direct-to-Consumer^5^ (DTC), saliva-based genotyping platform. Once generated, such a generic profile of over 600,000 SNPs could assist medical practitioners in diagnosing this as well as thousands of other diseases on demand, virtually genotyping cost free.

## Motivation

The conception of this paper was directly stimulated by a desire to respond to a harsh critique^6^ of two widely publicised recent studies^7,8^ of polygenic risk score (GRS) models for disease risk prediction. This critique by Wald & Old, provocatively titled “Illusion of Polygenic Disease Risk Prediction” is concluded with a pessimistic statement “*To our knowledge, no genome-wide polygenic score meets* [a sufficiently strong association] *requirement, and none is likely to do so with polygenic scores that emerge in the future*” in order “*to be seriously considered as a possible screening test*”.

We share concerns about ability of GWASs assembling thousands of unrelated individuals to demonstrate convincing translational relevance of generated results, especially when those are gauged against well-established clinical tests or some genetic tests discovered in family studies. However, we do not share such total pessimism at all. Actually, we see numerous exciting novel opportunities, in particular, resulting from combing genomics with diverse complementary testing techniques into improved novel diagnostic protocols. This does not seem to be accounted for by Wald & Old^6^ and thus motivated this articulation of our perspective on that matter.

The ideal genomic test for case-control GWAS should split data into two complementary subsets, one with very high, the other with very low penetrance of disease cases. In practice this is hardly ever possible, since genomes determine only susceptibility for disease development, while the actual onset of disease depends also on other factors, e.g. environmental triggers, which are typically not accessible during GWAS analysis. In some practical applications this can be compensated by combining genomics with independent clinical tests which effectively detect history of those exposures. However, one may reasonably expect that in some extreme genotype configurations the susceptibility is so high that sufficient, relatively weak triggers are almost certainly met by the carriers who become diseased. Similarly, on the opposite protective extreme, the “necessary-for-disease-development” genomic variants could be missing and so these people will be almost certainly disease free. Detection of such extreme genomic configurations has obvious practical implications^1–4^, even if it could be of relevance for a small fraction of the population only. An obvious motivating example here is the case of deleterious mutations in the BRCA1 or BRCA2 genes^9,10^: around 5-10% of breast cancer (BC) cases in women are attributed to some harmful mutations in those genes with estimated 5 fold increased risk (or penetrance) to 80% for breast cancer (by the age 90)^10^. There are hundreds of known mutations in those genes, of which only some are known to be deleterious with approximately equal attribution of the risk between both genes. No one will question these days that a genetic test detecting some of those deleterious mutations in BRCA1 gene is of prime medical concern, even if it could be of relevance for far less than 2.5-5% of possible BC cases, as it could a trigger need for some drastic preventive actions by some of the unfortunate carriers^11^. Against this background, we shall proceed to re-analyse five CD risk models for the detection of highly deleterious and highly protective genetically determined strata, in particular.

## GWAS Results

We use GWAS data for CD composed of four different ethnic cohorts: British (UK), Dutch (NL), Finish (FIN) and Italian (IT), comprised altogether of 3796 cases and 8154 controls (see Suppl. Table 1). Genotyping data were accessed from https://www.ebi.ac.uk/ega/studies/EGAS00000000057 under accession number EGAD00010000286 and originally used in the series of papers ^12 1,2^ reporting the main risk models used here. The largest UK cohort (1849 cases and 4936 controls) was used for discovery (training) where applicable. The combined remaining 3 cohorts, *FIN* ∪ *IT* ∪ *NL*, were used for independent testing (1947 cases & 3218 controls). We consider 5 different models of risk stratification as follows:

i. GRS_228_ which uses 228 SNPs selected by the a penalised regression applied to the UK cohort^1^;
ii. HDQ_15_ based on specific configurations of 4 Human Leucocyte Antigen (HLA) haplotypes imputed to data using HIBAG_HLA algorithm ^13^ and splitting data into 15 disjoint CD risk strata ordered according to the risk assessed in the UK cohort ^2^; see Suppl. Table 4;
iii. HDQ_17_ – based on calls of 6 directly measured HLA SNPs and developed by application of machine learning and human expertise to UK-cohort data^2^; see Suppl. Table 5;
iv. TD – a HLA-based CD risk model splitting data into 5 different expert determined risk categories developed independently of this GWAS data by Tye-Din, *et al*,^4^;
v. ROM - another HLA-based CD risk model splitting data into 3 different, expert-determined risk categories developed by Romanos, *et al*.^3^.

Note that that GRS_228_ using 228 SNPs generates stratification with over 11,000 different values in our data, which is complex and ‘noisy’ in comparison to a handful of level-based *parsimonious* stratifications from any of remaining four models. In Figure 1 we show five Receiver Operating Characteristic (ROC) curves for those risk models with the Area Under the Curve (AUC), a typical metric used to quantify classification performance, displayed in the legend. All curves, except for ROM model, look very similar, especially in the extreme ends: the ‘deleterious’ left-hand-side-bottom corner marked as a pink rectangle and the ‘protective’ right-hand-side upper corner, marked in green. In the zoomed inserts we start seeing some differences between individual ROC curves, though they are still hard to interpret. We need to examine the shape of the extreme ends of these ROC curves, focusing especially on the steepness at the curve onset as well as its flatness at the top. To that end, in Figure 2.A we plot a *precision-recall* curve in terminology of information retrieval[ref] or, in the epidemiological terms, the *penetrance* versus *sensitivity*, which is the fraction of diseased people in the sub-population of carriers of genotypes falling into the tail of the distribution versus the rate of detecting cases (y-coordinate in Figure 1). For the protective corner in Figure 2.B we plot penetrance vs. specificity, which is the rate of detecting disease-free controls (x-coordinate in Figure 1). The penetrance is estimated from the independent test data on the combined Finish, Italian and Dutch cohorts under prevalence *K*_*CD*_ = 1%, which is an estimated prevalence of CD in the European population^14^ as well as in Australia^4^ and USA^15,16^. In Figure 3 we show statistical significance (p-values) for all those models obtained from the Fisher exact test: we clearly see that all results for the tails larger than 1.5% are statistically very significant.

**Figure 1:**
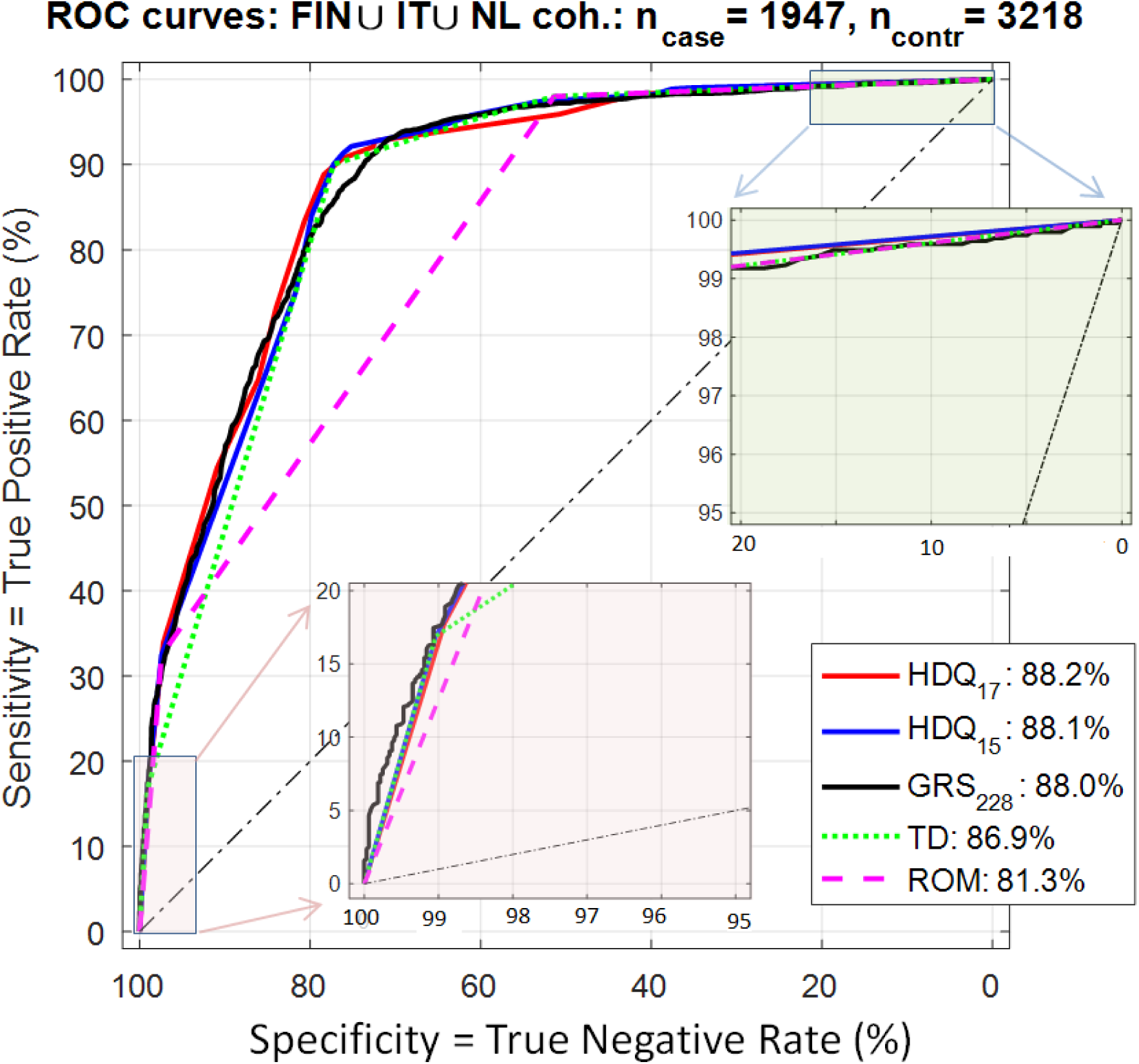
ROC Curves for five different risk models tested on the independent test data (the combined three cohorts Finish (FIN), Dutch (NL) and Italian (IT)). The extreme deleterious (South-West corner) and protective (North-East corner) stratification regions are marked as pink and blue rectangles, respectively, with two inserts presenting a higher resolution views. All curves, with an exception of TD model, look close to each other with hardly any difference noticeable by visual inspection. Note that the AUCs, shown in the legend, are very close to each other and, especially for the three top models, do not differentiate between these models in a meaningful way.

**Figure 2:**
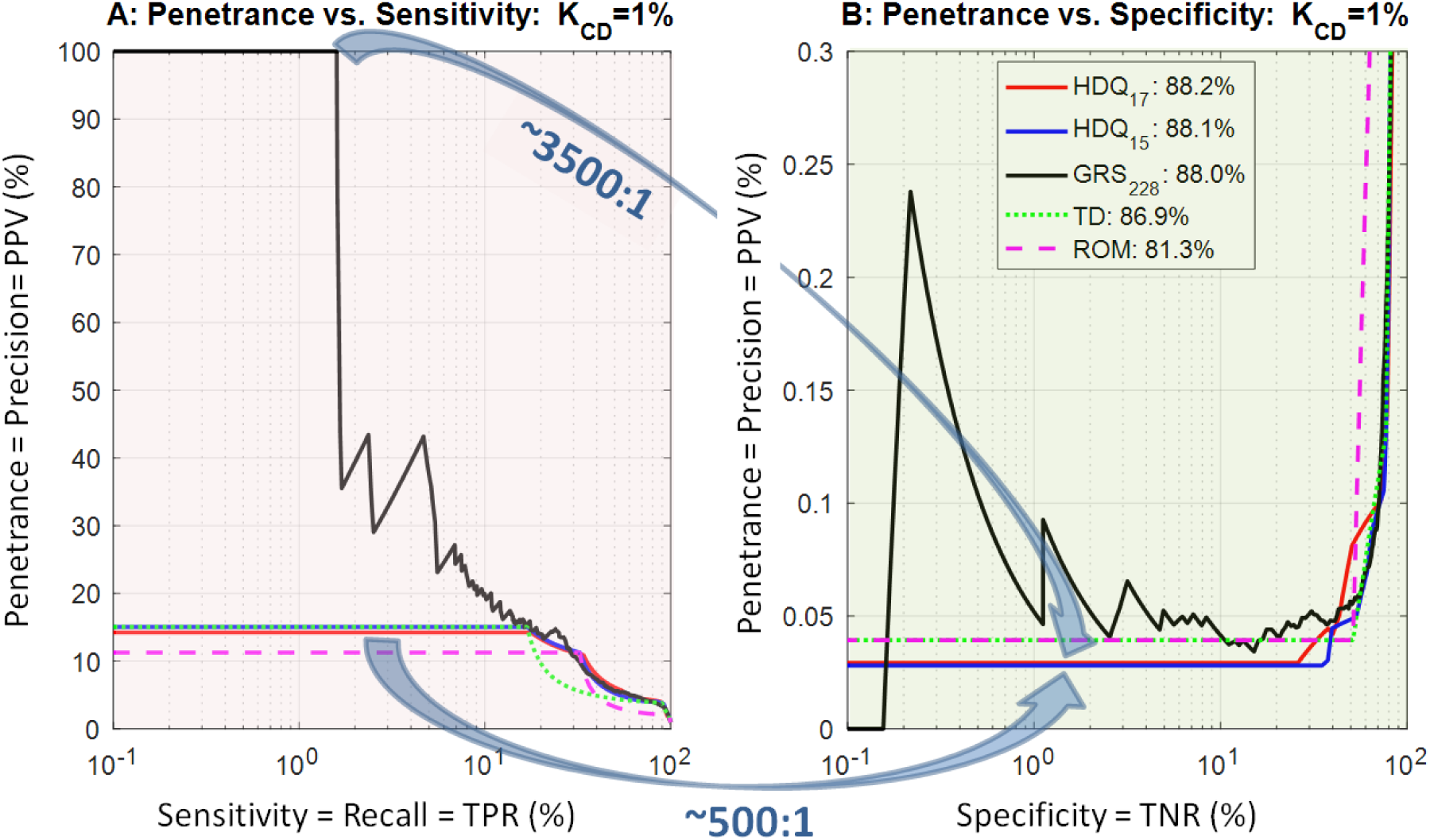
Plots penetrance versus sensitivity and specificity for the independent test data (FIN ∪ IT ∪ NL cohorts combined). (**A**) The penetrance vs. sensitivity or, equivalently, the precision vs. recall, or the *positive predictive value* (PPV) vs. *true positive rate* (TPR), etc. (**B**) The penetrance vs. specificity, or PPV vs. true negative rate (TNR), etc. We assume prevalence K_CD_=1%, an estimated population penetrance of coeliac disease in the European population^14^ as well as in Australia^4^ and USA. Two panels are designed to show unfolding of singularities at the deleterious and protective corners of the ROC curves shown as pink and blue rectangles in Figure 1, respectively. We show results of testing on the combined three off-training cohorts (Finish, Dutch and Italian). Note the highly desired significant drops in prevalence between the most deleterious strata in left panel and the most protective strata in the right panel with impressive ratios, 3500: 1 and 500:1, in particular. Note also that all parsimonious risk models, i.e. other than GRS_228_, have very stable tails (horizontal tail lines) extending to 20-30% in the x-axis dimension. On the other hand, the plots for GRS_228_ are relatively noisy, which is expected, as they reflect contributions from over 200 independent terms. Furthermore, the parsimonious models clearly outperform GRS_228_ at the protective extremes (right panel) which, somewhat indirectly, reflects a relatively good knowledge of link between coeliac disease and relatively common HLA variants. However, at the deleterious corner (left panel) GRS_228_ clearly outperforms all remaining models reaching ultimate penetrance 100% for the extreme stratum encompassing ∼1.5% of CD-cases (p-value < **10**^−**9**^, see Figure 3). None of these remarkable features and differences are apparent if analysis is reduced to comparisons of AUC values or a superficial visual inspection or ROC curves in Figure 1.

**Figure 3:**
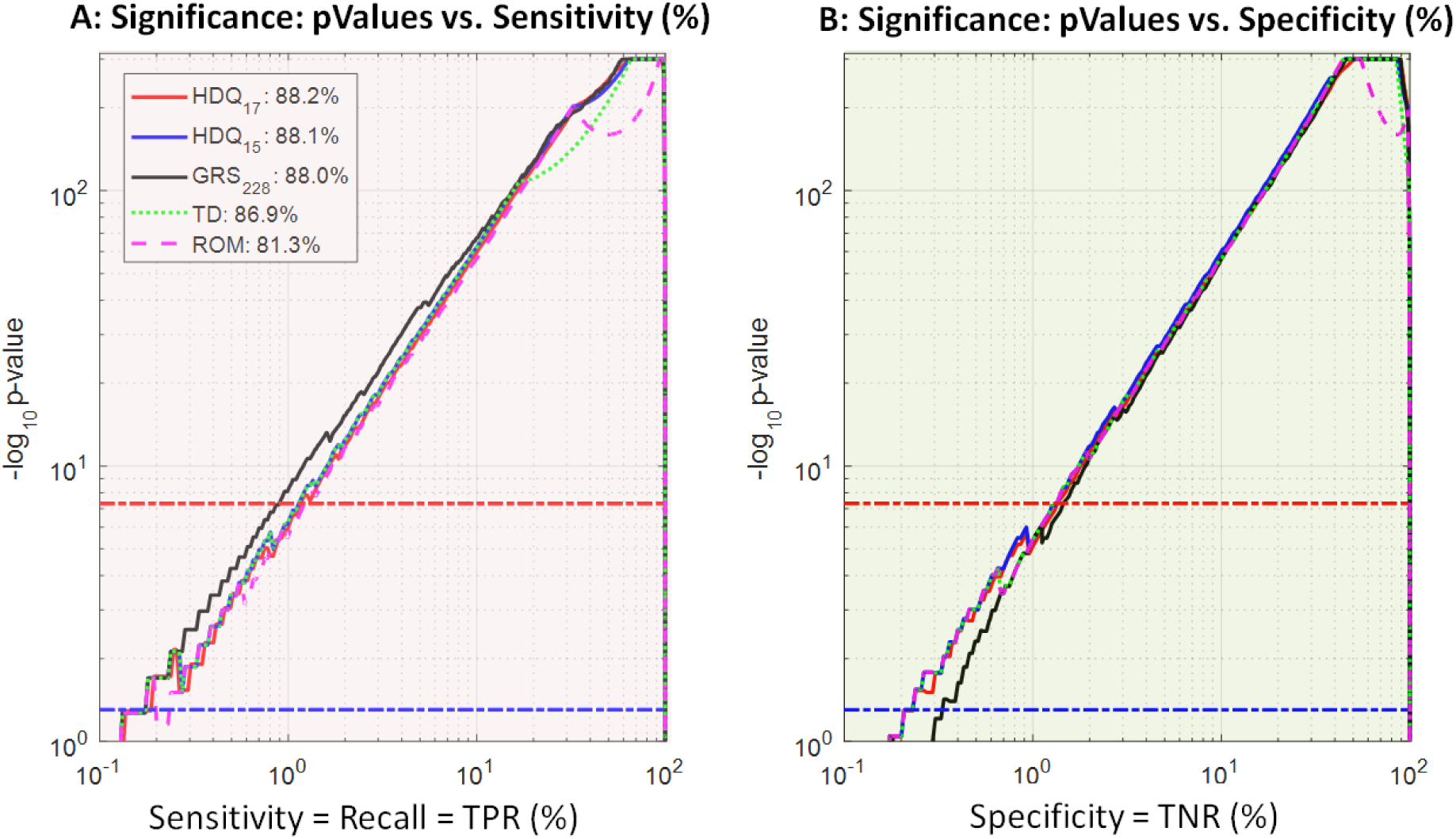
Significance for the extreme tail strata according to the Fisher exact test. The nominal **p**_**value**_ = **0. 05** and the standard genome wide **p**_**value**_ = **5** × **10**^−**8**^ significance levels are marked with blue and red horizontal chain lines. Note that in this independent test all extreme tail-strata of size ≳ **1. 5**% become very significant, with p-values ≲ **10**^−**8**^. Note that even the nominal level 0.05 of significance (blue chain line) is appropriate as there are no penalties for multiple choices involved here.

The key observations underpinning our analysis can be summarised as follows. Firstly, in Figure 1 we observe a characteristic ‘kink’ at the sensitivity level ∼90% splitting data into a ‘deleterious’ part containing 90% of cases and the ‘protective’ part containing remaining ∼10% of cases. Secondly, some “tiny” differences in ROC curves in Figure 1 between GRS_228_ and other model are actually linked to very significant differences in the performance as shown in Figure 2. In particular, in Figure 2A we see that the GRS_228_ model at the extreme ‘deleterious’ tail (for sensitivity ≈ 1.5%) conveys the ultimate risk, the penetrance 100%. This is 6 times larger than the highest risk for any of the remaining four models. These specific genotype configurations picked by GRS_228_ (present in 9FIN + 5IT + 17NL + 18UK samples) have a potential to progress aetiological knowledge of CD and should be investigated further. Moreover, we observe the penetrance ∼40% for GRS_228_ high risk tail containing up to 5% of CD cases, again warranting a follow up investigation.

On the opposite, ‘protective end’, GRS_228_ performs noticeably worse than any remaining four parsimonious models. It is represented here by a relatively noisy (black) curve, presumably affected by a noise injected by multiple, redundant probes. At this extreme, by using HDQ_15_ or HDQ_17_ risk models we can select subpopulations encompassing ∼30% controls, with very low disease risk (penetrance ∼0.03%), which corresponds to a drop by a ratio ∼3500:1 with respect to the top penetrance segment of 1.5% for GRS_228_. If you directly use the parsimonious CD-risk stratification based on the HDQ_17_ model (red curve), then we identify the top risk segment encompassing 18% cases with penetrance >14%, which is ∼500 times higher than the penetrance <0.03% observed for the bottom 25% of risk distribution according to that model.

### Commodity Genotyping

As it has been mentioned above the genetic testing required by our risk models is very simple: for comparison, the risk models used by Khera et al. ^7^ and Inouye et al. ^8^ are relying on millions of SNPs and other covariates. The simple genotyping required by our models can be delivered at a very low cost leveraging popular DTC genotyping platforms. For example, Illumina’s GSA microarray^5^ utilised by AncestryDNA or 23andme, already contains 4 out of 6 SNPs used by HDQ_17_ model and 61 SNPs used by *GRS*_228_; other required SNPs could be appended or replaced by appropriate proxies. Independently, ROM, TD and HDQ_15_ models can be readily implemented on such platforms by using additional HLA imputation algorithms. As for the future, the genotyping of the whole population is becoming a practical possibility. At the current cost of AU$99 per person in Australia (MyHeritageDNA™) it will cost AU$2.5Billion to genotype every of 25Million Australians, which is 1.4% of the estimated annual health expenditure in Australia 2016–17 (AU$180.7Billions; reference^17^, page 5). Once such profiles are generated, they could be utilised by their owners as cost-free, life-time resources for the assessment of life style choices and of risks for thousands of diseases (it is estimated that there are ∼8,000 heritable conditions of interest which new-borns should be tested^18^). Such sharing will reduce cost of genotyping to a negligible expense per diagnosis, or to 0, if a person has already acquired such a popular profile for some other reasons. Note that AncestryDNA kit is currently, June 2019, advertised by Amazon for USA at US$ 99.

### Translational Implications

Now we translate the above observations into three specific clinical scenarios with some estimates of costs and benefits (see Methods and Supplementary Materials for more detailed explanations). Given that only 30% to 40% of CD cases are currently diagnosed, improving the diagnosis of CD is now recognized as an important goal for clinicians [6] and application of genomic risk prediction lowering costs as outlined below can contribute towards practical realisation of this goal.

#### Detecting highly penetrant rare variants in general population

For argument sake, we estimate that in Australia the subpopulation with CD risk of 100% estimated using GRS_228_ model (Figure 2.A) is around 4,000 people (approximately 25Million / 6,700). If a simple saliva-based GRS_228_-like risk score had been used to detect this vulnerability early in their life, then these people would have been saved from coeliac suffering since birth, by adopting an appropriate diet. In case of USA, with 327Million citizen, this translates to ∼49,000 people. Note that these cohorts although relatively small, are actually larger than the total numbers of sufferers from many other malicious multifactorial conditions we do care about such as Primary Sclerosing Cholangitis (PSC), the prime indication for liver transplant, with prevalence in Europe^19^ ranging from 1/446,000-1/6,170. The searching for such rare variants would become very economical if multiple (hundreds) of diseases could be evaluated in parallel using a single DNA profile.

#### Detecting CD in the symptomatic subpopulation

The estimated CD prevalence in the cohort showing CD suggestive symptoms^1,16,20^ is 3%. Using prevalence ratio as a guide, we have estimated that 33% (=1%/3%) of the population displays CD-symptoms. This means that at some point ∼8.3Million of Australians need to be considered for lower bowel endoscopy, ‘the gold standard’ CD-confirmatory procedure. The total cost of that procedure would be AU$8.3 Billion (@AU$1000 / person^1,4^), with efficiency 1:33, i.e. of 1 CD-hit per 33 endoscopies. However, using prioritisation by HDQ_17_ risk model (or any of other 4 models but TD), we expect to detect 50% and 90% of CD cases with hit rates increased to 1: 5.8 and 1:9.2 by using only ∼725,000 and ∼2,029,000 endoscopies, respectively (see Supplementary Table 6 and Supplementary Figures 6). In comparison to random application of the procedure to 50% and 90% of this cohort this will save 3,442,000 and 5,426,000 endoscopies costing AU$3.442Billion and AU$5.426Billion, respectively.

By extrapolating these savings to the population of 327Million of US citizens, where cost of endoscopies (in US$) are very similar to Australia, the estimated saving would be of ∼US$45.0 Billion and ∼US$ 70.9 Billion for detection of 50% and 90% of CD-cases, respectively.

The genotyping overheads are ≤ AU$0.83 Billion and ≤ US$ 10.9 Billion, respectively, which is the cost of genotyping de novo of each person in the cohort at $100 /person. This, in the worst case, will make only 24% and 15% of estimated above savings on endoscopies, for targeted 50% and 90% CD-detection rates, respectively.

#### Detection of CD for the first-degree relatives of CD sufferers

We estimate that the cohort of first-degree relatives of CD sufferers (CD_FDR_) in Australia has approximately 0.25 Million ≈ 25Million × 1% × 33% × 3 people (assuming prevalence in the general population 1% with only 33% CD-cases diagnosed and 3 CD-undiagnosed first-degree relatives per known CD case). In this cohort, with prevalence^1,21,22^ of 10%, we expect 1 CD case detected per 10 endoscopies in random screening. However, if we select ∼30% of this cohort with the highest risk according to HDQ_17_ risk model we should detect ∼90% CD-cases with average hit rate 1 per 3.3 endoscopies. This would save >151,000 endoscopies and AU$ 151Million in comparison with screening of 90% of CD_FDR_ uniformly. This saving scaled to the USA will amount of US$1.975Billion. Note that the detection of the remaining 10% CD cases in CD_FDR_ requires significantly increased efforts, generating in excess of 30 endoscopies per single successful detection on average (see Supplementary Table 7 and Figure 7). The genotyping overheads for this scenario are ≤ AU$25 Million and ≤ US$ 327Million, respectively, which is below 17% of the estimated savings on endoscopies.

Interestingly, the estimated efficiency of 1 CD hit per 3.3 endoscopies discussed above is much higher than for an alternative, CD-dedicated blood based HLA typing test, which^1^ ‘at 10% CD prevalence […] would generate over five unnecessary endoscopies per correct endoscopy’. Thus this once-off test offers significantly lower hit rate (1:6) at a higher cost, namely, at AU$120/sample and typically US $150/sample or greater in USA^1^.

## Discussion

In simple terms, the savings in the last two scenarios leverage ability of genomics risk models to concentrate 90% of CD cases in 30% of a cohort translating to a 3-fold increase in efficiency of the expensive confirmatory procedures. The main cost saving comes from sacrificing the detection of the remaining 10% of cases, which would require scanning of the remaining 70% of the cohort with efficiency 21 times lower (21= (90%/30%): (10%/70%)). One may expect that similar results could be achieved using alternative, more established diagnostic techniques, e.g. serological and histological assessments^1,4,23^, but genomic risk assessment shows here translational potential and should be seriously considered as an alternative or a complement to those techniques. For instance, under our modelling assumptions, if the confirmatory endoscopy test is directed to the stratum of the symptomatic cohort containing 50% of the highest HDQ_17_-risk cases (Suppl. Table 6, row 4), we expect to detect 125,000 CD cases, which is more than 75,000-100,000 CD cases (30%-40%) currently diagnosed^6–8,24,25^ in Australia. This detection can be achieved using 125,000 × 5.8 ≈ 725,000 endoscopies, which is ≈ 8.7% of the symptomatic cohort and only ∼2.9% of the Australian population. Note that in this case the endoscopy hit rate of over 1:5.8 is almost twice better than the disease frequency among siblings of CD-cases.

Obviously, this could be achieved over years, gradually as patients with suggestive symptoms turn for assessment. (Such a delay could be, for instance, due to an earlier asymptomatic period of disease or to a development of disease in adulthood ^22,23,26–29^, which could also lead to repeated assessments of some individuals, increasing effective size of the symptomatic cohort, etc.)

Finally, note that all five CD risk models considered here comfortably exceed formal benchmarks used or reported in paper ^7–8,24,25^ and used by Wald and Old as foundation of their pessimistic paper^6^ (see Supplementary Tables 2 and 3 and a discussion following them). Thus, this plainly falsifies Wald and Old’s the statement^6^ that no genome-wide polygenic risk score models are likely to meet their formal standards of association accuracy^23,24^. However, we still concede that none of these models fulfils classical expectations for a medical screening test standing on its own, and also that none is able of explaining the majority of CD heritability^4^, which is often expected of genomic risk modelling. But as it has been argued here and elsewhere^1,2^ that does not preclude those models from producing potentially highly useful practical results which is an ultimate goal in this game. In particular, we have demonstrated above that relatively moderate risk prediction improvements, of order 3 - 4 times over cohort average, could lead to substantial translational gains in suitable clinical context as long as the genomic risk assessment could be delivered at affordable cost. However, this is a statement of technical capability. The final success will require to overcome a number of additional hurdles, such as the issue of privacy, reliability of genotyping tests and a demonstration of benefits in rigorous clinical trials among the other issues.

## Conclusions

On this note we re-iterate our opinion that the pilot results, such as those outlined above for CD, have enormous translational potential and should be seriously considered for developing/incorporation into novel disease screening tests along more established clinical techniques. The compact genotyping facilitating such risk models allows for accommodation of profiles for hundreds of other disorders using a single commodity, DTC-like, genotyping platforms, which will promote ubiquity and minimise diagnostic costs.

Finally, we should stress that success of such deployments depends also on successful development of supporting datamining techniques and dedicated software tools for data driven detection of rare variant strata conveying statistically significant extreme associations, discovery of compact (parsimonious) risk models and for incorporation of interactions among genomic and non-genomic factors for further improvements in model compactness and accuracy.

## Acknowledgements

The author acknowledges European Genome-phenome Archive for access to data to undertake our study (Project EGAD0001000028).

Many thanks to Justin Bedo, Richard Campbell, Michael Erlichster, Benjamin Goudey, Miroslaw Kapuscinski, Kerrie, Mark & Waclaw Kowalczyk and Nicolas Wong for assistance in preparation of this manuscripts.

## Methods

All five risk models used in the paper are exactly the same as in ^2^. All risk models were calibrated on UK cohort, the remaining 3 cohorts were used for an independent test. In summary:

1. **GRS**_**228**_: We recall, this risk model uses 228 SNPs selected by the penalised regression applied to the UK cohort (Abraham et al. 2014; see Supplementary Methods for more detail). An explicit text file describing the model, grs.txt, and a procedure for the generating risk scores from it are available at http://dx.doi.org/10.6084/m9.figshare.154193. This procedure generates 11,863 different scores for our data, hence for a simplicity of data processing and plotting, these scores were rounded to 500 levels uniformly distributed between the lowest and the highest value.
2. **ROM, TD, HDQ**_**15**_: For these 3 models required 4-digit HLA-DQA1 and HLA-DQB1 genotypes for each sample were imputed using the R package HIBAG (HLA Genotype Imputation with Attribute Bagging) ^13^. Based on the specific combination of those haplotype each sample was allocated to one of the 3 for ROM ^3^, of 6 for TD ^4^ and of 15 for HDQ_15_ ^2^ subsets (strata). For each stratum the risk of disease was represented by the ratio of CD-cases to the fraction of controls falling into it, which is a data estimate of the positive likelihood ratio (LR+) (sometimes also referred to as *the positive likelihood ratio* (*PLR*) of the *odds ratio with reference to the population odds* (*OR*). The explicit formulae follow for completeness and the numerical values are given in Supplementary Tables 4 & 5.

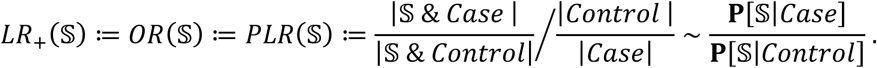

(Here 𝕊 denotes a stratum in question; | | denotes the cardinality and **P** stands for the probability and *case* / *control* denote the subset of cases and controls in the data.) In our binary classification analysis (bases on ROC curves analysis) only the order of strata is really accounted for (but not the explicit values of scoring function, i.e. *LR*_+_ in this case). We have used the ordering on UK cohort to that end. This ordering agrees perfectly with the risk order allocated in the original papers introducing ROM and TD models.
3. **HDQ**_**17**_: This model introduced in ^2^ stratifies data into 17 categories according to the allele calls for 6 specific SNPs, see Suppl. Table 5. As before we have used the *LR*_+_(𝕊) values for UK-cohort in Suppl. Table 5 in order to rank the CD-risk for those strata.

Given the prevalence *K*_*CD*_ of the disease in the population and a genomic stratum 𝕊 we estimated the *penetrance f*(𝕊), i.e. the probability of being diseased while a carrier of genotype in 𝕊, also known as the *positive predictive value* (*PPV*):

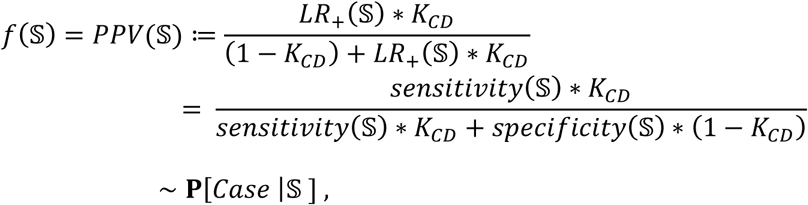

Where

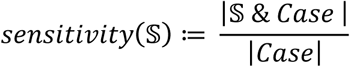

And

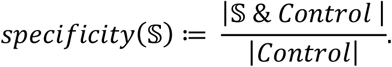

Note that in the main results of this paper we have exclusively analysed strata 𝕊 defined by the extreme deleterious or extreme protective tails of the risk distributions (see Figure 2, the Supplementary Figures 2, 4 and Supplementary Tables 4 & 5).

The inverse of the penetrance provides a dataset-based estimate of the number of tests (endoscopies) which will be required to detect a single CD-case in the genomic stratum 𝕊. It has been used in generation of Supplementary Tables 6&7 and Figures 5&6 on which the discussion of the Translational Implications in the paper was based.

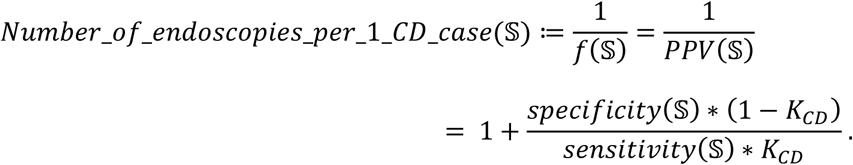

Each stratum 𝕊 splits our case-control data into a 2×2 contingency table. The classical Fisher Exact test was used to assess significance of such partitions against a null hypothesis that 𝕊 was selected by random drawing of |𝕊| samples from the data. For the extreme protective and extreme deleterious tails of the risk distributions such p-values are plotted in Figure 3 and Supplementary Figure 3.

