## Supplementary Materials for "Predictive and Aetiological Potential of Polygenic Strata with Extreme and Moderate Disease Risks Predictions: *Case Study of Coeliac Disease GWAS*"

Adam Kowalczyk [1-8],....

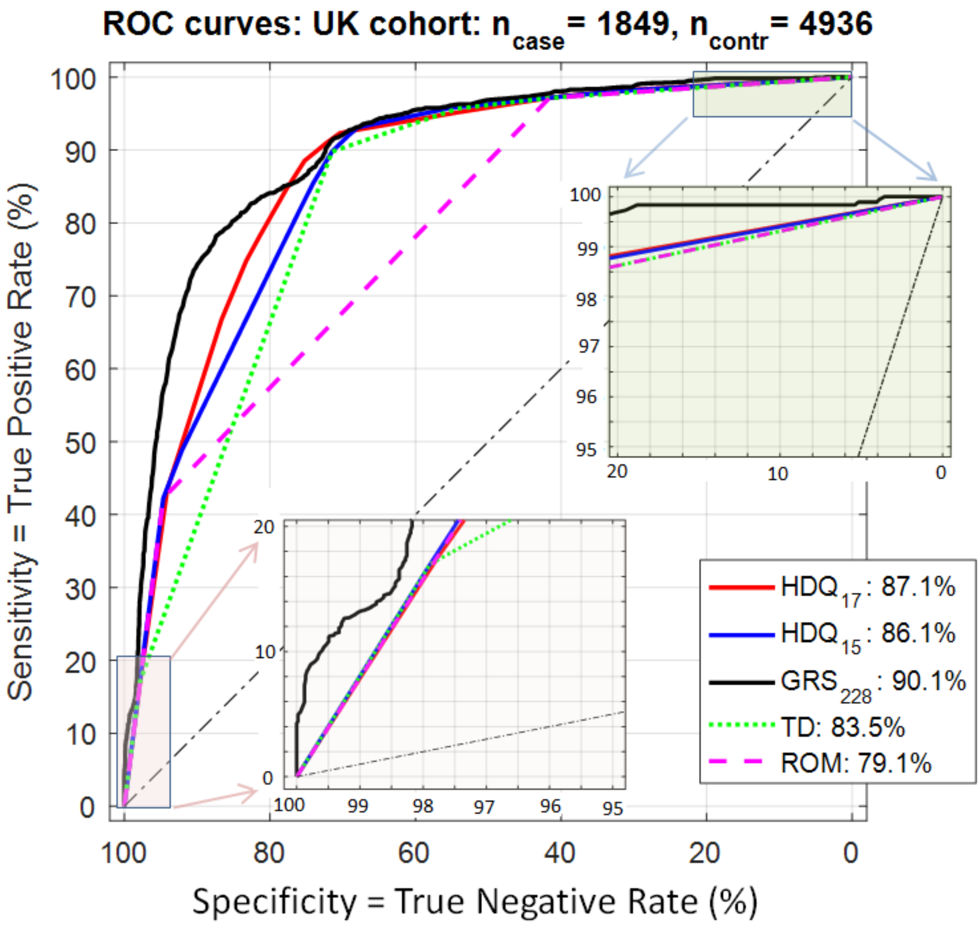


**Supplementary Figure 1: ROC Curves for five different risk models on the discovery data (UK cohort).**

In contrast to the independent test in Figure 1 of the paper, the $GRS_{228}$ model is clearly outperforming all the other models. However, one should be cautious here as these differences involve an entanglement of some ethnic differences with an overfitting to the training data.

[1]Centre for Neural Engineering, [2] Department of Computing and Information Systems & [3]Centre for Epidemiology and Biostatistics, The University of Melbourne, Melbourne, Victoria, Australia; [4]Diversity Arrays Technology Pty. Ltd., Canberra, Australia; [5]Genomic Biomarkers Systems, Pty. Ltd., Melbourne, Australia; [6]Centre for Eye Research Australia, Melbourne, Australia; [7] Institute for Applied Ecology, University of Canberra, Canberra Australia; [8]Faculty of Mathematics and Information Science, Warsaw University of Technology, Poland;

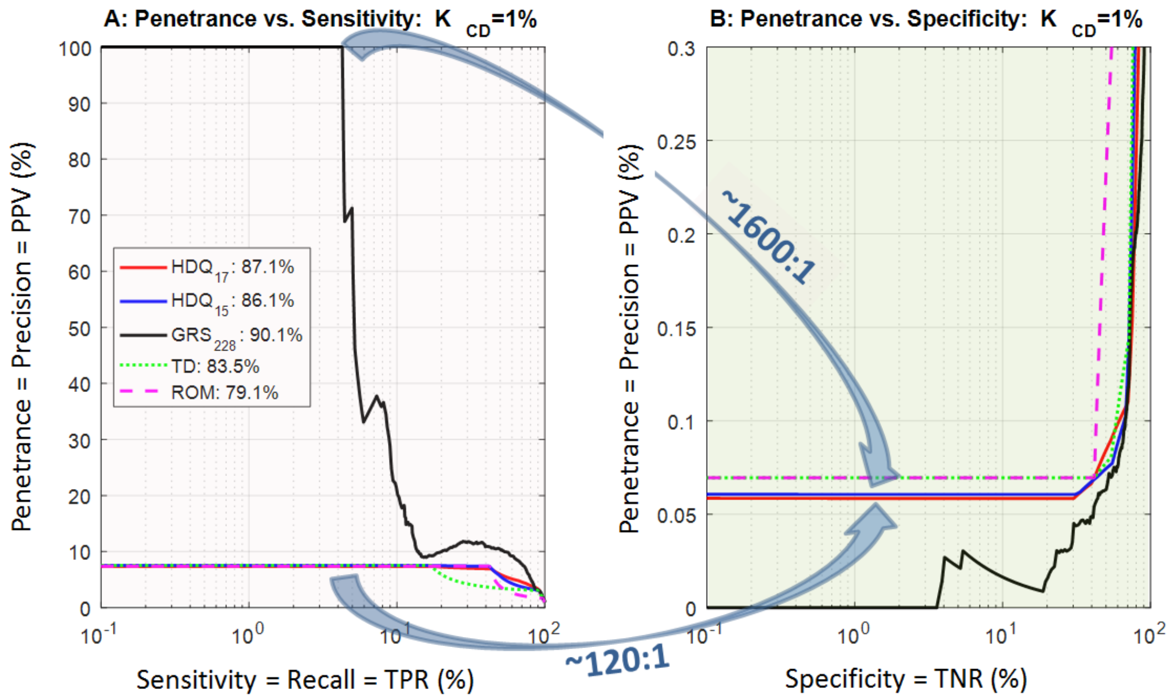


**Supplementary Figure 2: Plots for the discovery data (UK cohort): (A) the penetrance vs. sensitivity or, equivalently, the precision vs. recall and (B) the penetrance vs. specificity. We assume prevalence $K_{CD}$=1% (an estimate of the population penetrance of coeliac disease in Australia or USA).**

Note that the highest risk stratum for $GRS_{228}$ model (penetrance 100%) is here over 4% in size compared to the 1.5% in non-UK cohorts in Figure 2. However, the penetrance in the protective extreme (right panel) is also higher, ~0.06% compared to ~0.03% in Figure 2. These differences could be most likely explained by the differences in distribution of HLA alleles between UK and other European ethnicities, which are known to exist. Obviously, a follow-up investigation of these differences is warranted here.

.

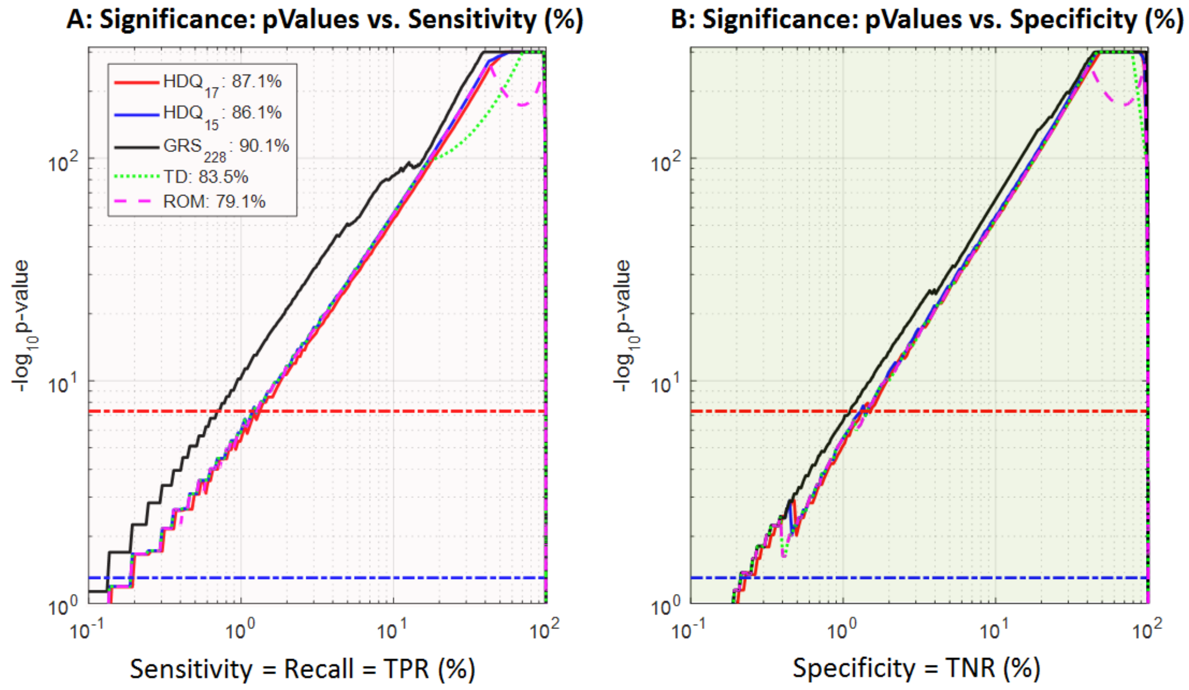


**Supplementary Figure 3: Significance for the discovery data (UK cohort) for the extreme strata shown in Supplementary Figure 2 according to the Fisher exact test.**

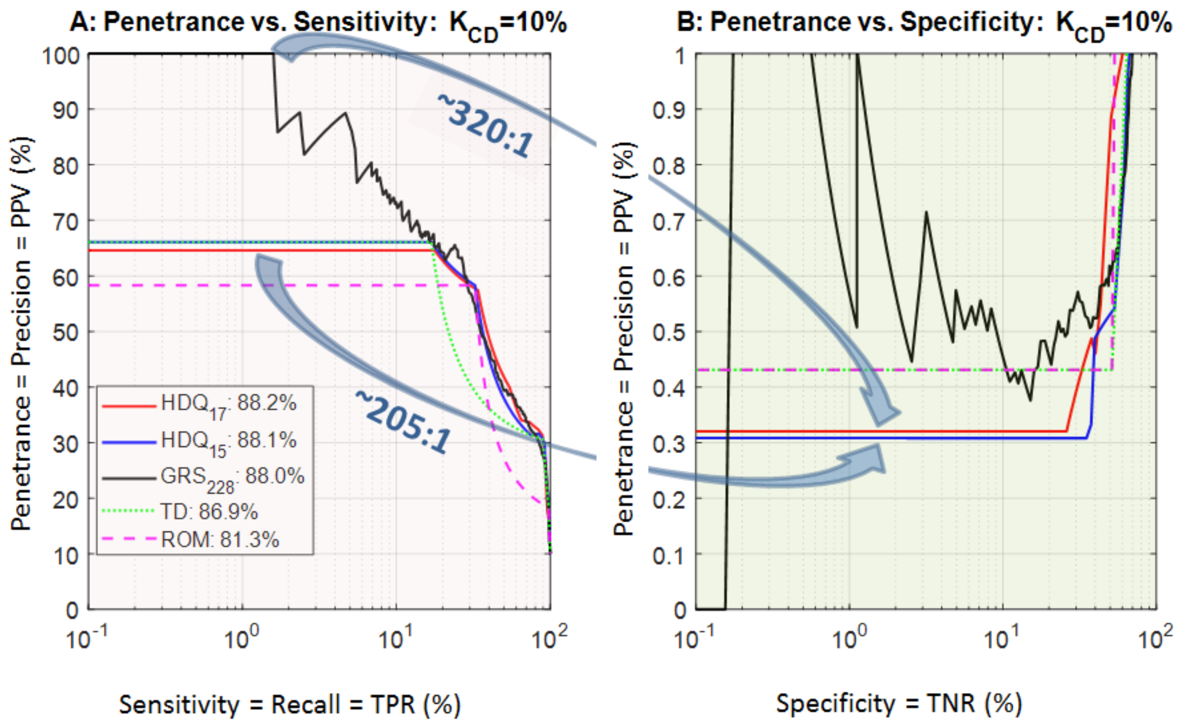


**Supplementary Figure 4: Penetrance for extreme tails test on 3 non-UK cohorts FIN ∪ IT ∪ NL scaled to $K_{CD}$=10% - the estimated prevalence among first degree relatives (FDR) of CD-sufferers** (Bourgey et al. 2007; Abraham et al. 2014; Rubio-Tapia et al. 2008)**. (A) the penetrance vs. sensitivity or, the precision vs. recall and (B) penetrance vs. specificity.**

Note that here the highest risk strata for $HDQ_{15}$ & $HDQ_{17}$ cover ~17% of CD cases with impressive penetrance >65%. Obviously, the penetrance in the protective extreme covering ~33% controls (right panel) is also higher than before, ~0.3% versus ~0.03% in Figure 2. Still we observe an impressive penetrance drop ratios ~200 :1 between the highest and the lowest risk strata for $HDQ_{15}$ and $HDQ_{17}$. Note that in this case each of the three, the highest two CD-risk strata for either $HDQ_{15}$ of $HDQ_{17}$ and the top stratum for ROM, cover ~34% of CD-cases, with penetrance ~58%. Thus again, hypothetically, a simple saliva-based test can potentially preselect a subpopulation of FDRs, with a risk increased over 5-fold in comparison to FDR-average. Thus, in principle at least, a simple saliva–based genetic test on GSA array can save around 33% of FDRs from colonoscopy, or at least postpone need of such an invasive procedure until later age, in the case of juveniles.

| Population | SNP dataset | Post QC SNPs | Cases | Controls |
|---|---|---|---|---|
| **UK** | Illumina Hap550 | 528,969 | 1,849 | 4,936 |
| **FIN** | Illumina Hap550 | 528,969 | 647 | 1,829 |
| **NL** | Illumina Hap550 | 528,969 | 803 | 846 |
| **IT** | Illumina Hap550 | 528,969 | 497 | 543 |
| Total | | | 3,796 | 8,154 |

saliva–based genetic test on GSA array can save around 33% of FDRs from colonoscopy, or at least

**Supplementary Table 1: Summary information on our cohorts for Coeliac Disease GWAS used for generation of three risk models (GRS228** (Abraham et al. 2014) **and $HDQ_{15}$ & $HDQ_{17}$** (Erlichster et al. 2019)**) discussed in the main paper.**

The data has been ben accessed from https://www.ebi.ac.uk/ega/studies/EGAS00000000057 under accession number EGAD00010000286 (UK2, FIN, IT, NL). It has been originally used in the paper (van Heel et al. 2007) as well as in a number of our papers in the past

| | Detection rates (=Sensitivity=TPR) | | | | |
|---|---|---|---|---|---|
| FPR | HDQ_17 | GRS_228 | TD | ROM | HDQ_15 |
| 1% | 16.4% | **17.6%** | 17.0% | 12.6% | 17.3% |
| 5% | **41.1%** | 39.9% | 30.5% | 35.9% | 38.9% |
| 10% | 56.41% | **56.45%** | 47.2% | 43.0% | 52.5% |
| 20% | **85.0%** | 81.5% | 80.8% | 57.3% | 83.2% |
| 25% | **91.2%** | 88.3% | 90.8% | 64.4% | 92.2% |

**Supplementary Table 2: The detection rates for selected false positive rates (FPR) for independent test on the combined 3 cohorts $\mathbf{FIN \cup IT \cup NL}$.**

Note that the *detection rate* used in ( Wald and Old 2019) is equivalent to sensitivity or true positive rates (TPR). The values for FPR=5% in the Table exceed significantly the values for CAD disease reported in (Khera et al. 2018; Inouye et al. 2018), namely, 15% and 13% respectively. These detection rates were criticized as too low for diagnostic purposes by (Wald and Old 2019).

**Supplementary Table 3: Odds ratio (OR) for independent test on the combined 3 cohorts** FIN ∪ IT ∪ NL. The results were obtained by interpolation of stratification according to the Supplementary Tables 3-6 of Erlichster et al. (2019), which for $HDQ_{15}$ and $HDQ_{17}$ models is explicitly repeated and expanded in the Supplementary Tables 4 & 5 below. Table 3.A is directly comparable with Table 3 of Khera et al. (2018). Here we estimate the x% of the score distribution by a segment x% of controls, which is a reasonable approximation of the population with a relatively low prevalence (1%) of CD and constitutes a lower, pessimistic, bound in comparison to the exact estimates.

A: $OR_{pop}$ - the relative odds: Ratio of odds in top x% vs remaining (100-x)%

| x% | **$HDQ_{17}$** | $GRS_{228}$ | TD | ROM | $HDQ_{15}$ |
|---|---|---|---|---|---|
| 20% | **22.7** | 17.6 | 16.8 | 5.4 | |
| 10% | 11.65 | **11.66** | 8.1 | 6.8 | |
| 5% | **13.3** | 12.6 | 8.3 | 10.6 | |
| 1% | 19.5 | **21.1** | 20.3 | 14.3 | |

B: Odds Ratios (RO): Ratio of odds in top x% top x% vs odds at bottom x%

| x% | **$HDQ_{17}$** | $GRS_{228}$ | TD | ROM | $HDQ_{15}$ |
|---|---|---|---|---|---|
| 20% | **147.5** | 99.1 | 103.7 | 73.5 | |
| 10% | **196.6** | 137.4 | 121.2 | 110.4 | |
| 5% | **289.4** | 155.4 | 156.3 | 184.3 | |
| 1% | **627.5** | 342.0 | 436.8 | 322.9 | |

Note that all odds ratios in Table A above exceed any equivalent x% entry in the Table 3 in Khera et al. (2018).

The first row in Table B effectively represents quintile relative odds according to (Wald, Hackshaw, and Frost 1999; Wald and Old 2019), which is defined as the ratio of odds between people in the 1st and the 5th quintiles (20%) of risk distribution. All those values exceed significantly the value 50 cited as an acceptable minimum (Wald and Morris 2011). Further, these numbers are significantly higher than the comparable quintile hazard ratio of 4.17 for CAD achieved by a metaGRS-model using over million DNA variants (Inouye et al. 2018). This hazard ratio value was harshly criticised by (Wald and Old 2019) as far too low to be useful for diagnostic purposes.

**Supplementary Table 4: Outline of the $HDQ_{15}$ risk model** (Erlichster et al. 2019). Each category was defined by a pair of HLA alleles as specified in the second column the Table. The genotypes imputed to the data using HIBAG_HLA algorithm (Zheng et al. 2014). All entries with missing calls were omitted. The rows are sorted in the decreasing order of the positive likelihood ratio (LR+) for the discovery UK cohort and this risk ordering was used in test on the non-UK cohorts (FIN ∪ IT ∪ NL). Note the discrepancies in the order of LR+ values between the discovery and the test data, e.g. in rows 3, 4 & 5; 8 & 9, etc.

| | | Discovery: UK | | | | | Independent test: FIN ∪ IT ∪ NL | | | | |
|---|---|---|---|---|---|---|---|---|---|---|---|
| # | Genotype, HLA alleles | LR+ | Case % | Control % | pentr $K_{CD}$=1% | pentr $K_{CD}$=10% | LR+ | Case % | Control % | pentr $K_{CD}$=1% | pentr $K_{CD}$=10% |
| 1 | DQ2.5/DQ2.5 | 8.01 | 17.2% | 2.2% | 7.5% | 44.5% | 16.19 | 17.1% | 1.1% | 14.1% | 64.3% |
| 2 | DQ2.5/DQ2.2 | 7.25 | 25.0% | 3.5% | 6.8% | 42.0% | 9.33 | 15.7% | 1.7% | 8.6% | 50.9% |
| 3 | DQ2.5/DQ8 | 2.49 | 6.4% | 2.6% | 2.5% | 19.9% | 2.49 | 6.6% | 2.6% | 2.5% | 21.7% |
| 4 | DQ2.2/DQ7 | 2.06 | 32.9% | 15.9% | 2.0% | 17.1% | 2.71 | 36.4% | 13.4% | 2.7% | 23.1% |
| 5 | DQ8/DQ8 | 1.87 | 3.6% | 1.9% | 1.9% | 15.8% | 4.00 | 8.1% | 2.0% | 3.9% | 30.8% |
| 6 | DQ2.5/DQX | 1.58 | 3.7% | 2.3% | 1.6% | 13.6% | 2.39 | 5.3% | 2.2% | 2.4% | 21.0% |
| 7 | DQ8/DQ2.2 | 0.95 | 2.3% | 2.4% | 1.0% | 8.7% | 1.04 | 1.3% | 1.2% | 1.0% | 10.4% |
| 8 | DQ8/DQ7 | 0.90 | 1.1% | 1.2% | 0.9% | 8.3% | 0.74 | 0.8% | 1.1% | 0.7% | 7.6% |
| 9 | DQ2.5/DQ7 | 0.24 | 0.3% | 1.4% | 0.2% | 2.4% | 4.34 | 0.7% | 0.2% | 4.2% | 32.5% |
| 10 | DQ2.2/DQX | 0.21 | 2.5% | 11.9% | 0.2% | 2.0% | 0.23 | 3.3% | 14.3% | 0.2% | 2.5% |
| 11 | DQ8/DQX | 0.16 | 2.0% | 12.1% | 0.2% | 1.6% | 0.30 | 2.4% | 8.2% | 0.3% | 3.2% |
| 12 | DQ7/DQ7 | 0.11 | 0.9% | 8.7% | 0.1% | 1.1% | 0.07 | 0.9% | 12.8% | 0.1% | 0.8% |
| 13 | DQ7/DQX | 0.09 | 0.1% | 0.6% | 0.1% | 0.9% | 0.00 | 0.0% | 2.2% | 0.0% | 0.0% |
| 14 | DQ2.2/DQ2.2 | 0.06 | 0.1% | 1.7% | 0.1% | 0.6% | 0.21 | 0.4% | 2.0% | 0.2% | 2.3% |
| 15 | DQX/DQX | 0.06 | 1.9% | 31.6% | 0.1% | 0.6% | 0.03 | 1.0% | 35.0% | 0.0% | 0.3% |

**Supplementary Table 5: Outline of the $HDQ_{17}$ risk model** (Erlichster et al. 2019). Each category was defined by a combination of alleles of six defining SNPs as indicated. All instances with missing calls for any of those 6 SNPs were neglected. The rows are sorted in the decreasing order of the positive likelihood ratio (LR+) for the discovery UK cohort and this risk ordering was used in test on the non-UK cohorts (FIN ∪ IT ∪ NL). Note the discrepancies in the order of LR+ values between the discovery and the test data, e.g. in row 10.

| | Defining SNP_alleles | | | | | | | Discovery: UK | | | | | Independent test: FIN ∪ IT ∪ NL | | | | |
|---|---|---|---|---|---|---|---|---|---|---|---|---|---|---|---|---|---|
| # | rs3129763_T | rs2187668_T | rs2856705_A | rs9275312_G | rs9357152_C | rs9275572_C | ~HLA genotype | LR+ | case% | control% | pentr $K_{CD}$=1% | pentr $K_{CD}$=10% | LR+ | case% | control% | pentr $K_{CD}$=1% | pentr $K_{CD}$=10% |
| 1 | <=2 | 2 | 0 | 0 | - | - | DQ2.5/DQ2.5 | 7.85 | 17.8% | 2.3% | 7.34% | 44.0% | 16.44 | 17.7% | 1.1% | 14.24% | 62.2% |
| 2 | <=1 | 1 | 1 | 0 | - | - | DQ2.5/DQ2.2 | 7.07 | 25.7% | 3.6% | 6.67% | 41.4% | 9.44 | 16.2% | 1.7% | 8.71% | 48.6% |
| 3 | <=1 | 1 | 0 | 0 | 0 | 2 | DQ2.5/DQ6.2_2 | 3.13 | 23.3% | 7.5% | 3.06% | 23.8% | 3.26 | 20.6% | 6.3% | 3.19% | 24.6% |
| 4 | <=1 | 1 | 0 | 1 | - | - | DQ2.5/DQ8 | 2.38 | 8.0% | 3.4% | 2.35% | 19.2% | 4.07 | 8.4% | 2.1% | 3.95% | 28.9% |
| 5 | 1 | 0 | 1 | 0 | - | - | DQ2.2/DQ7 | 1.83 | 3.7% | 2.0% | 1.82% | 15.5% | 3.06 | 10.1% | 3.3% | 3.00% | 23.4% |
| 6 | <=1 | 1 | 0 | 0 | 0 | <=1 | DQ2.5/DQ6.2_1 | 1.75 | 6.3% | 3.6% | 1.74% | 14.9% | 2.14 | 10.3% | 4.8% | 2.12% | 17.6% |
| 7 | 2 | 1 | 0 | 0 | - | - | DQ2.5/DQ7 | 1.56 | 3.8% | 2.4% | 1.55% | 13.5% | 2.42 | 5.5% | 2.3% | 2.39% | 19.5% |
| 8 | 0 | 0 | 1 | 1 | - | - | DQ2.2/DQ8 | 0.81 | 2.4% | 3.0% | 0.81% | 7.5% | 0.86 | 1.9% | 2.3% | 0.86% | 7.9% |
| 9 | 0 | 0 | 0 | 2 | - | - | DQ8/DQ8 | 0.73 | 1.4% | 1.9% | 0.73% | 6.8% | 0.38 | 1.3% | 3.6% | 0.38% | 3.6% |
| 10 | 0 | 0 | 2 | 0 | - | - | DQ2.2/DQ2.2 | 0.20 | 0.3% | 1.4% | 0.20% | 2.0% | 4.07 | 0.6% | 0.2% | 3.95% | 28.9% |
| 11 | 0 | 0 | 0 | 1 | - | - | DQ8/DQX | 0.17 | 2.6% | 15.0% | 0.17% | 1.7% | 0.26 | 1.9% | 7.4% | 0.26% | 2.5% |
| 12 | 0 | 0 | 1 | 0 | - | - | DQ2.2/DQX | 0.16 | 1.9% | 12.2% | 0.16% | 1.5% | 0.15 | 3.2% | 21.7% | 0.15% | 1.4% |
| 13 | 1 | 0 | 0 | 1 | - | - | DQ8/DQ7 | 0.13 | 0.3% | 2.2% | 0.13% | 1.3% | 0.15 | 0.5% | 3.3% | 0.15% | 1.4% |
| 14 | 2 | 0 | 0 | 0 | - | - | DQ7/DQ7 | 0.09 | 0.1% | 0.6% | 0.09% | 0.9% | 0.08 | 0.9% | 11.8% | 0.08% | 0.8% |
| 15 | 1 | 0 | 0 | 0 | - | - | DQ7/DQX | 0.09 | 0.8% | 8.8% | 0.09% | 0.9% | 0.03 | 0.8% | 26.0% | 0.03% | 0.3% |
| 16 | 0 | 0 | 0 | 0 | - | - | DQX/DQX | 0.06 | 1.8% | 30.2% | 0.06% | 0.6% | 0.00 | 0.0% | 2.3% | 0.00% | 0.0% |
| 17 | <=1 | 1 | 0 | 0 | 1 | 0 | DQ2.5/DQ7.3 | 0.00 | 0.0% | 0.0% | 0.00% | 0.0% | 0.00 | 0.0% | 0.1% | 0.00% | 0.0% |

| Detected CD-cases | | Number of endoscopies per 1 CD-detection | | | | | Saved endoscopy costs ($ Million) | | | | |
|---|---|---|---|---|---|---|---|---|---|---|---|
| % | # | $HDQ_{17}$ | $HDQ_{15}$ | $GRS_{228}$ | TD | ROM | $HDQ_{17}$ | $HDQ_{15}$ | $GRS_{228}$ | TD | ROM |
| 1.5% | 3,750 | 3.0 | 2.8 | 1 | 2.8 | 3.6 | 114 | 114 | 121 | 114 | 112 |
| 18% | 45,000 | 3.0 | 2.9 | 2.9 | 2.9 | 3.6 | 1,366 | 1,370 | 1,371 | 1,368 | 1,339 |
| 33% | 82,500 | 3.6 | 3.8 | 4.2 | 4.3 | 3.7 | 2,450 | 2,440 | 2,404 | 2,399 | 2,441 |
| 50% | 125,000 | 5.8 | 6.3 | 6.4 | 5.7 | 7.2 | 3,442 | 3,378 | 3,363 | 3,454 | 3,272 |
| 90% | 225,000 | 9.2 | 9.2 | 10.4 | 9.2 | 15.4 | 5,426 | 5,437 | 5,151 | 5,439 | 4,042 |

**Supplementary Table 6: Estimates of savings in cost and number of endoscopies in detection of CD in Australian cohort of people with suggestive CD-symptoms ($CD_{Sympt}$ cohort). All risk models used are as in the independent test on $\mathrm{FIN} \cup \mathrm{IT} \cup \mathrm{NL}$ -cohort (Figure 1 of the paper)**

The size of $CD_{Sympt}$ is estimates as 8.33 Million (= 25M Australians * 1% CD prevalence / 3% prevalence in $CD_{Sympt}$ , see Abraham et al. 2014; Catassi et al. 2007; Hin et al. 2011). As before we use an estimate of $1000 for low bowel endoscopy cost and assumed ROC curves in Figure 1 of the paper for the risk modelling (see also Figure 5 below). The cost of colonoscopy for testing whole $CD_{Sympt}$ @ $1000 per person is AUD $8.33 Billion; the cost of de novo genotyping @ $100 per person is $ 833 Million. This difference will imply clear cost savings and justify genotyping based prioritisation even if we apply colonoscopy aiming only at the top 50% of CD cases (row 4 in Table). This should detect more than the estimated of 30%-40% (75,000 - 100,000) of CD-cases being currently diagnosed in Australia (Anderson 2011; Catassi et al. 2007; Dubé et al. 2005; Anderson et al. 2013). Note that improving the diagnosis of CD is now recognized as an important goal for clinicians (Thompson 2005) and genomic risk prediction can clearly contribute toward realisation of this goal.

The number of endoscopies per 1 detected CD-case is estimated as the inverse of penetrance $1/f(\mathbb{S}) \ = 1/PPV(\mathbb{S})$ , see Methods.

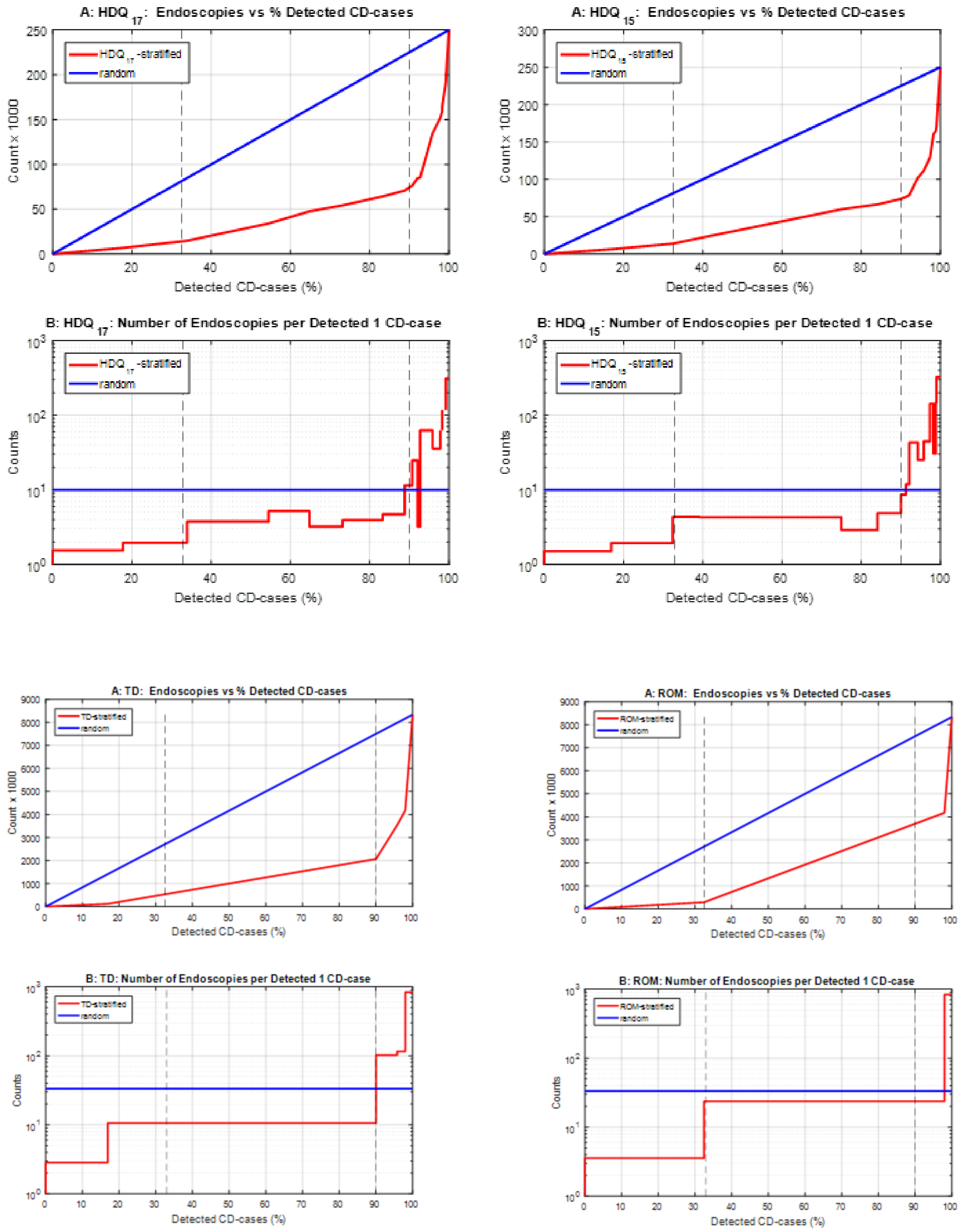


**Supplementary Figure 5: Estimated numbers of small bowel endoscopy of for scanning $CD_{Sympt}$ cohort of 8.33 Million people using preselection according to $HDQ_{17}$, $GRS_{228}$, TD & ROM CD-risk models.** The number of endoscopies per 1 detected CD-case is estimated as the inverse of penetrance $\mathbf{1}/\boldsymbol{f}(\mathbb{S}) = \mathbf{1}/\boldsymbol{PPV}(\mathbb{S})$ , see Methods. All risk models used here are as in the independent test on $\mathbf{FIN} \cup \mathbf{IT} \cup \mathbf{NL}$ -cohort, with the ROC-curves as in Figure 1 of the paper.

Note the substantial increase in the number of endoscopies per detected CD case in the detection of the last 10% of CD-cases (e.g. a clear jump from <10 to >100 in the case of TD model).

| Detected new CD-cases | | Number of endoscopies per 1 CD-detection | | | | | Saved endoscopy costs ($Million) | | | | |
|---|---|---|---|---|---|---|---|---|---|---|---|
| % | Number | $HDQ_{17}$ | $HDQ_{15}$ | $GRS_{228}$ | TD | ROM | $HDQ_{17}$ | $HDQ_{15}$ | $GRS_{228}$ | TD | ROM |
| 33% | 8,250 | 1.75 | 1.79 | 1.9 | 2.6 | 1.85 | 68 | 68 | 67 | 61 | 67 |
| 50% | 12,500 | 2.39 | 2.63 | 2.47 | 2.94 | 3.68 | 95 | 92 | 94 | 88 | 79 |
| 90% | 22,500 | 3.29 | 3.27 | 3.64 | 3.27 | 5.3 | 151 | 151 | 143 | 151 | 106 |

**Supplementary Table 7: Estimates of savings in cost and number of endoscopies in detection of CD in the high-risk cohort of first-degree relatives ($CD_{FDR}$ cohort).**

The size of $CD_{FDR}$ is estimates as 250,000 (= 25M Australians × 1% CD prevalence × 0.25 diagnosed cases rate × 4 undiagnosed FDRs per diagnosed case) containing 25,000 CD-cases assuming 10% cohort prevalence (Bourgey et al. 2007; Abraham et al. 2014; Rubio-Tapia et al. 2008). For simplicity we have used an estimate $1000 for a low bowel endoscopy cost- “the ‘gold standard’ confirmatory test” for CD following (see discussion in Abraham et al. 2014). Our estimates are based on ROC curves in Figure 1 of the paper (test on the independent data, see Supplementary Figures 6 below). The de novo genotyping costs of this $CD_{FDR}$ cohort @ $100 per person is $25 Million, hence even at the worst-case scenario, when this cost is incurred in full, there will be substantial savings if endoscopies are guided by genomic risk predictions. However, more important here is the avoidance of unnecessary endoscopies, due to inconvenience of procedures and the danger of complications. Thus, an innocuous genotyping test based on saliva facilitating such savings is a very attractive option, especially, in the case of juveniles. Interestingly, it is estimated that if an alternative, non-invasive blood based test, HLA typing were used as a guide for further investigations, at 10% CD prevalence it would generate over five unnecessary endoscopies per successful detection, and this test in known to be even less accurate for children below 4 years old (Abraham et al. 2014). Five endoscopies per CD-case is substantially higher than the estimated numbers of endoscopies for any method but ROM in the above Table.

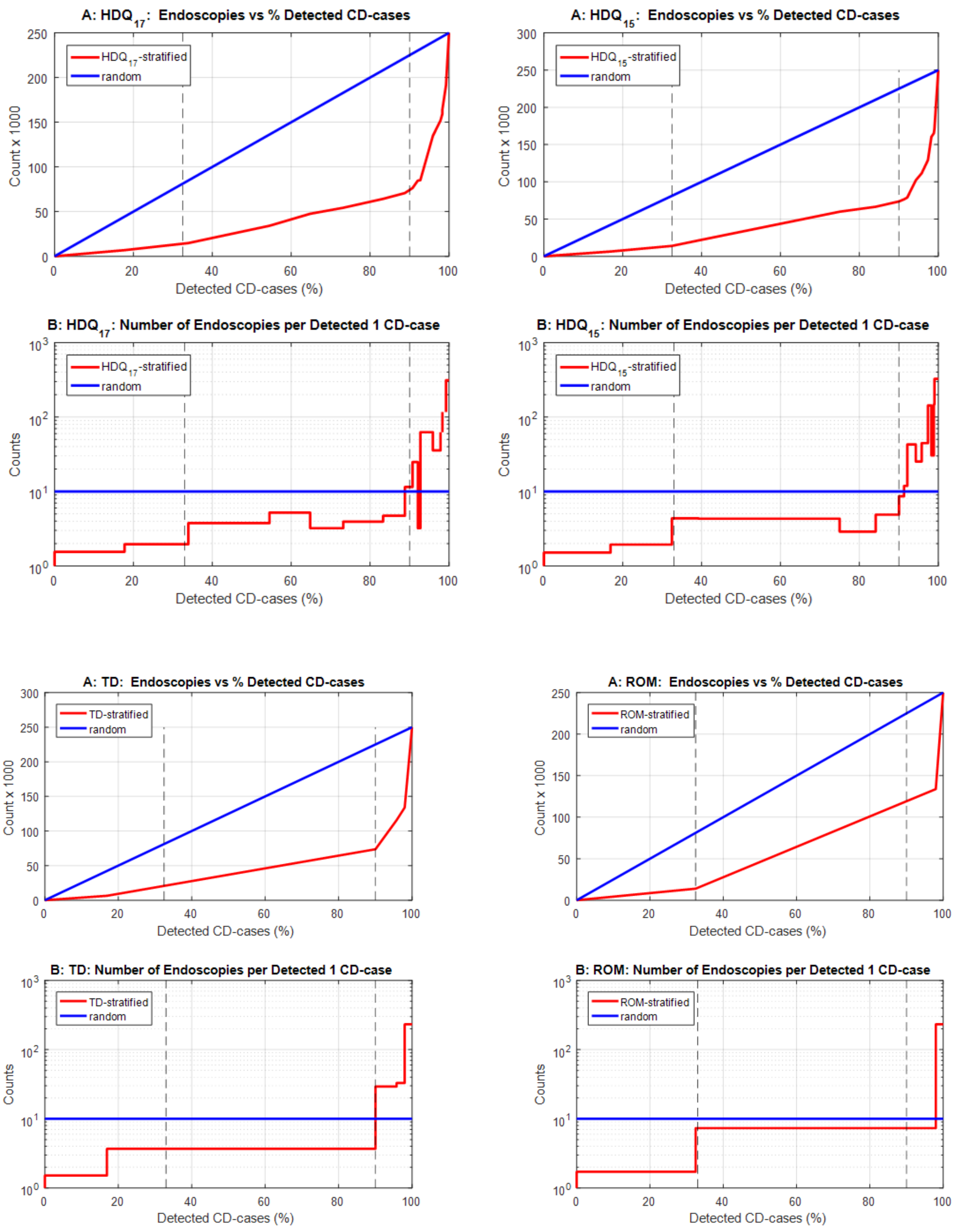


**Supplementary Figure 6: Estimated numbers of small bowel endoscopy for scanning 250,000 first degree relatives of CD sufferers (CD$_{FRS}$ cohort) using preselection according to HDQ$_{17}$, HDQ$_{15}$, TD and ROM CD-risk models.** The number of endoscopies per 1 detected CD-case is estimated as the inverse of penetrance $\mathbf{1}/\boldsymbol{f}(\mathbb{S}) = \mathbf{1}/\boldsymbol{PPV}(\mathbb{S})$, see Methods. All risk models are as in the independent test on $\mathbf{FIN} \cup \mathbf{IT} \cup \mathbf{NL}$ -cohort, with the ROC-curves as in Figure 1 of the paper.

Note the substantial increase in the number of endoscopies pre detected CD case in the detection of the last 10% of CD-cases (e.g. a clear jump from <3.7 to >29 in the case of TD model.

https://doi.org/https://doi.org/10.1136/bmj.319.7224.1562.

Wald, Nicholas J., and Joan K. Morris. 2011. "Assessing Risk Factors as Potential Screening Tests: A Simple Assessment Tool." *Arch Intern Med.* 171 (4): 286–91. https://doi.org/10.1001/archinternmed.2010.378.

Wald, Nicholas J, and Robert Old. 2019. "The Illusion of Polygenic Disease Risk Prediction." *GENETICS in MEDICINE* 0 (0): 1–3. https://doi.org/10.1038/s41436-018-0418-5.

Zheng, X, J Shen, C Cox, J C Wakefield, M G Ehm, M R Nelson, and B S Weir. 2014. "HIBAG--HLA Genotype Imputation with Attribute Bagging." *The Pharmacogenomics Journal* 14 (2): 192–200. https://doi.org/10.1038/tpj.2013.18.
